# Exploring vulnerable proteins in the progression of head and neck squamous cell carcinoma

**DOI:** 10.64898/2026.08.07.743269

**Authors:** Avantika Agrawal, Swapnil Kumar, Vaibhav Vindal

**Author notes:** Corresponding author: Dr. Vaibhav Vindal Department of Biotechnology & Bioinformatics, School of Life Sciences, University of Hyderabad, Gachibowli, Hyderabad, 500046, India., Email ID –.

## Abstract

A protein whose removal or deletion causes significant disruption or collapse of a protein-protein interaction (PPI) network is referred to as a vulnerable protein. Such proteins may serve as valuable therapeutic or diagnostic targets in disease-associated networks. In this study, two PPI networks were constructed, one for HPV-positive and the other for HPV-negative head and neck squamous cell carcinoma (HNSCC), and the vulnerable proteins of these networks were identified by the node deletion approach. After analyzing the networks, 27 unique vulnerable proteins in HPV-positive and 72 unique vulnerable proteins in HPV-negative HNSCC were identified. Among them, one HPV-positive and seven HPV-negative HNSCC vulnerable proteins were further chosen by integrating multi-omics data. To exploit the vulnerabilities of these proteins, candidate synthetic lethal (SL) partners were predicted whose inhibition may selectively impair tumor survival. Subsequently, drug-gene interaction analysis was performed to identify inhibitors targeting the SL partners of these vulnerable proteins. Notably, in HPV-positive HNSCC, *TOP2A, CHEK1*, and *CHEK2* genes were identified as SL partners of *TTN*, and their inhibitors were already clinically approved. While in HPV-negative HNSCC, *ADA* and *MMP19* were identified as an SL partner of *LMO7*; *TMEM45B, CDH3*, and *ELF3* genes were identified as an SL partner of *CGN*; and *ZNF433* was identified as an SL partner of *FLNC*. However, *MMP19, ZNF433*, and *TMEM45B* inhibitors were not reported. Thus, these vulnerable proteins, including their SL partners, provide novel avenues to explore and develop more efficient and precise therapeutic and diagnostic strategies.

## 1. Introduction

Head and neck squamous cell carcinoma (HNSCC) is a heterogeneous disease that arises in the squamous cells of the upper aero-digestive tract. As per the GLOBOCAN 2022 report, in India, around 2,39,817 new cases and 1,33,046 deaths were reported in both males and females with HNSCC (Bray *et al*., 2024). The risk factor of HNSCC includes excessive intake of tobacco and alcohol, use of cigarettes, and infection with human papillomavirus (HPV). HNSCC can be categorized into two groups based on the status of HPV infection: HPV-positive and HPV-negative HNSCC (Chandel *et al*., 2020). The HPV-negative HNSCC is caused by extravagant drinking of alcohol, and overconsumption of tobacco and smoking, while HPV-positive HNSCC is mainly caused by HPV. Several reports have shown that high-risk HPV infection is the major causative agent in the development of HNSCC (Svider *et al*., 2017). However, there has been a lack of systematic studies on HPV-positive and HPV-negative HNSCC, including underlying mechanisms and exploration of HPV-positive and HPV-negative specific candidate genes, despite ongoing research on HNSCC. Thus, it is important to understand the complex molecular mechanism and investigate the tumor-associated key proteins associated with HPV-positive and HPV-negative HNSCC.

Recently, network-based methods such as the protein-protein interaction (PPI) network have made remarkable achievements in understanding human diseases, including cancer. It is a complex disease that involves a large set of proteins interacting together to perform biological processes and molecular functions. Interactions between two or more proteins are determined experimentally using methods such as yeast two-hybrid (Schwikowski *et al*., 2000; Ito *et al*., 2000; Gavin *et al*., 2002), mass spectrometry (Ho *et al*., 2002; Pandey and Mann, 2000; Figeys, 2003), etc. Moreover, several databases such as APID (Alonso-López *et al*., 2019), HIPPIE (Alanis-Lobato *et al*., 2016), HuRI (Luck *et al*., 2020), and many more provide information about PPIs of various organisms. These databases provide both experimentally verified and predicted PPIs, and some of these databases are organism-specific. These PPIs data coupled with gene expression data of head and neck tumor tissues can be utilized for PPI network based analysis to understand and elucidate the mechanisms of HPV-positive and HPV-negative HNSCC in a more effective manner.

In this study, the differentially expressed genes (DEGs) of HPV-positive and HPV-negative HNSCC were mapped onto the known PPIs of human to construct PPI networks of HPV-positive and HPV-negative HNSCC. Further, Gene Ontology (GO) and Kyoto Encyclopedia of Genes and Genomes (KEGG) pathway analysis were performed to construct meaningful information from the HPV-positive and HPV-negative HNSCC proteins products of respective DEGs. This followed the calculation of the various topological properties of the network, like degree, betweenness, and clustering coefficient. Subsequently, the network vulnerability analysis was performed to find out the vulnerable proteins, protein pairs, and protein triplets in the network with the one-node deletion, two-node deletion, and three-node deletion approaches, respectively. Additionally, mutation, copy number variation, and synthetic lethal interactions of all vulnerable proteins were identified and analyzed for therapeutic relevance.

## 2. Materials and Methods

### 2.1 Gene expression data

Gene expression data of HNSCC were retrieved from TCGA using the TCGAbiolinks R package. The expression data contained a total of 498 tumor samples and 44 normal samples. Further, HPV-positive and HPV-negative HNSCC sample information of HNSCC was downloaded from the Broad Institute’s Firehose database (“https://gdac.broadinstitute.org/“). Based on the HPV-positive and HPV-negative HNSCC samples, 498 tumor samples were separated into 89 HPV-positive and 409 HPV-negative HNSCC samples.

### 2.2 PPI data downloaded from different sources and construction of PPI networks

The differentially expressed genes (DEGs) for both HPV-positive and HPV-negative HNSCC samples were collected from our previously published work (Agrawal *et al*., 2025). PPI data of HPV-positive and HPV-negative HNSCC were downloaded from various databases such as APID, HIPPIE, and HuRI. The DEGs of HPV-positive and HPV-negative HNSCC were mapped onto these PPI databases to obtain HPV-positive and HPV-negative HNSCC PPI datasets. The PPI datasets from the databases HPV-positive and HPV-negative HNSCC conditions were used to construct the networks of HPV-positive and HPV-negative HNSCC with the help of the igraph package (Csardi and Nepusz, 2006) in R. Further, degree distribution, along with degree-betweenness, and degree-clustering coefficient were explored in the HPV-positive and HPV-negative HNSCC PPI networks.

### 2.3 Functional and pathway enrichment analysis of proteins present in the network

To better understand the functions and pathways of the proteins present in the HPV-positive and HPV-negative HNSCC networks, Gene Ontology (GO) and KEGG pathway analysis were conducted by using the gProfiler2 package (Kolberg *et al*., 2020) in R. GO enrichment analysis of proteins present in the HPV-positive and HPV-negative HNSCC networks on the basis of biological processes, cellular components and molecular function while KEGG pathway analysis of proteins present in the HPV-positive and HPV-negative HNSCC networks helps us to identify the functions of the proteins that participated in the pathways.

### 2.4 Network vulnerability analysis

The vulnerability analysis of networks leads to the identification vulnerable proteins. The vulnerable proteins are those upon removal of which the structure and function of networks get altered or collapse. Vulnerable proteins (individual), protein pairs, and protein triplets were identified as key components of these networks by using NetVA package (Kumar *et al*., 2025) in R. To identify the vulnerable proteins, protein pairs, and protein triplets, each protein was knocked out of the original network randomly by a single-node deletion approach. The topological properties of the resultant network, such as average betweenness centrality, average closeness centrality, average eccentricity, average node connectivity, articulation point, and clustering coefficient, were calculated after the removal of each node to understand the vulnerability of the overall network structure. Thus, the robustness of the network was evaluated by the changes in the network structure after the removal of each node. To improve the quality and accuracy of the data and to ignore false positives, proteins present in at least two topological properties were considered. Proteins that were not reported in more than one topological property were removed. Second, interacting pairs of vulnerable proteins were knocked out from the network, using a two-node deletion approach, and various network topological properties of the resultant network were calculated. Based on any two properties, a list of vulnerable protein pairs was retrieved. In the same way, protein triplets (three proteins interacting) were knocked out of the network one by one using a three-node deletion approach, and various topological properties of the resultant networks were calculated. Based on at least two properties, vulnerable protein triplets were identified. The vulnerable proteins common in the single-node deletion, two-node deletion, and three-node deletion approaches were further used for analysis.

### 2.5 Mutations in vulnerable proteins

To understand the functional defectiveness of the vulnerable proteins of HPV-positive and HPV-negative HNSCC networks in-depth, somatic mutation and copy number variations analysis were conducted. Somatic mutation data were downloaded from TCGA using the TCGA biolinks Rpackage. The somatic mutation data consisted of various types of mutation such as missense mutation, nonsense mutation, silent, intron, and splice site. For vulnerable proteins, 3’Flank, 3’ UTR, intron and silent mutations were not considered, as these mutations do not change the structure of the proteins. In a similar way, gene-level copy number variation (CNV) data of TCGA processed with GISTIC2 were also retrieved from the UCSC Xena browser for further analysis.

For HNSCC, gene-level CNV data were classified based on the GISTIC2-derived copy number values. Genes with copy number values of -2 or -1 were categorized as deleted, those with values of +1 or +2 as amplified, and those with a value of 0 were considered copy number neutral. For vulnerable proteins, amplification and neutral regions were not considered because neutral genes have no change in copy number and are unlikely to influence gene dosage or function, while amplification genes are usually drivers of cancer.

### 2.6 Synthetic lethal partners of vulnerable proteins

Synthetic lethality is a phenomenon that describes the relationship in gene pairs within the cell. When one of the genes is missing or unavailable in a cell, the cell remains viable, while the loss of both genes leads to cell death (Fan *et al*., 2024). For example, The DNA damage pathway is controlled by the BRCA gene and the PARP gene. Somatic mutation in the BRCA gene can lead to the development of breast and ovarian cancer. In tumor cells with a BRCA mutation, the cell switched to its interacting partner, the PARP gene, and only the PARP gene would be active for the DNA damage pathway. The development of PARP inhibitor, niraparib, leads to blocking the DNA damage pathway and leads to cell lethality (Helleday, 2011).

Synthetic lethal partners of the previously identified vulnerable proteins were inferred by using SynLethDB (Wang *et al*., 2022), a well-known publicly available database. Only high-confidence SL pairs supported by experimental or computational evidence were retrieved. These interactions were further analyzed for therapeutic relevance by mapping known drug-gene associations from the DGIdb database (Cannon *et al*., 2024).

## 3. Result

### 3.1 PPI network construction

A total of 2276 and 2210 DEGs in HPV-positive and HPV-negative HNSCC, respectively, were collected from our previously published work (Agrawal *et al*., 2025). The interaction network consists of 2011 nodes (proteins) with 11895 edges (interactions) in HPV-positive HNSCC (Supplementary Figure F1) and 1894 nodes (proteins) with 9910 edges (interactions) in HPV-negative HNSCC (Supplementary Figure F2). To validate the HPV-positive and HPV-negative HNSCC networks, the degree distribution (Fig. 1), degree-betweenness (Fig. 2), and degree-clustering coefficient (Fig. 3) plots were visualised.

**Fig. 1:**
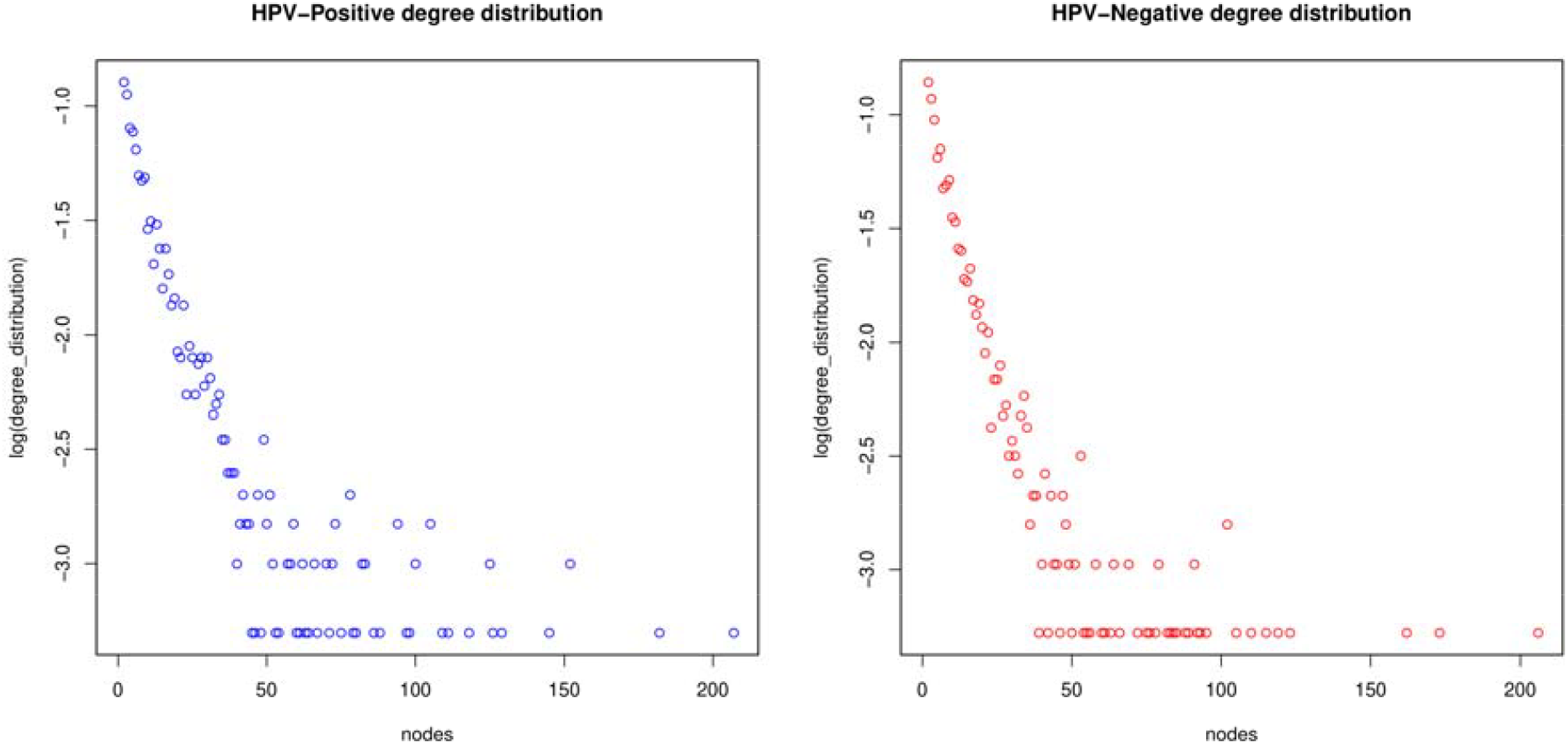
Degree distribution plot for HPV-positive (left-side) and HPV-negative (right-side) HNSCC networks.

**Fig. 2:**
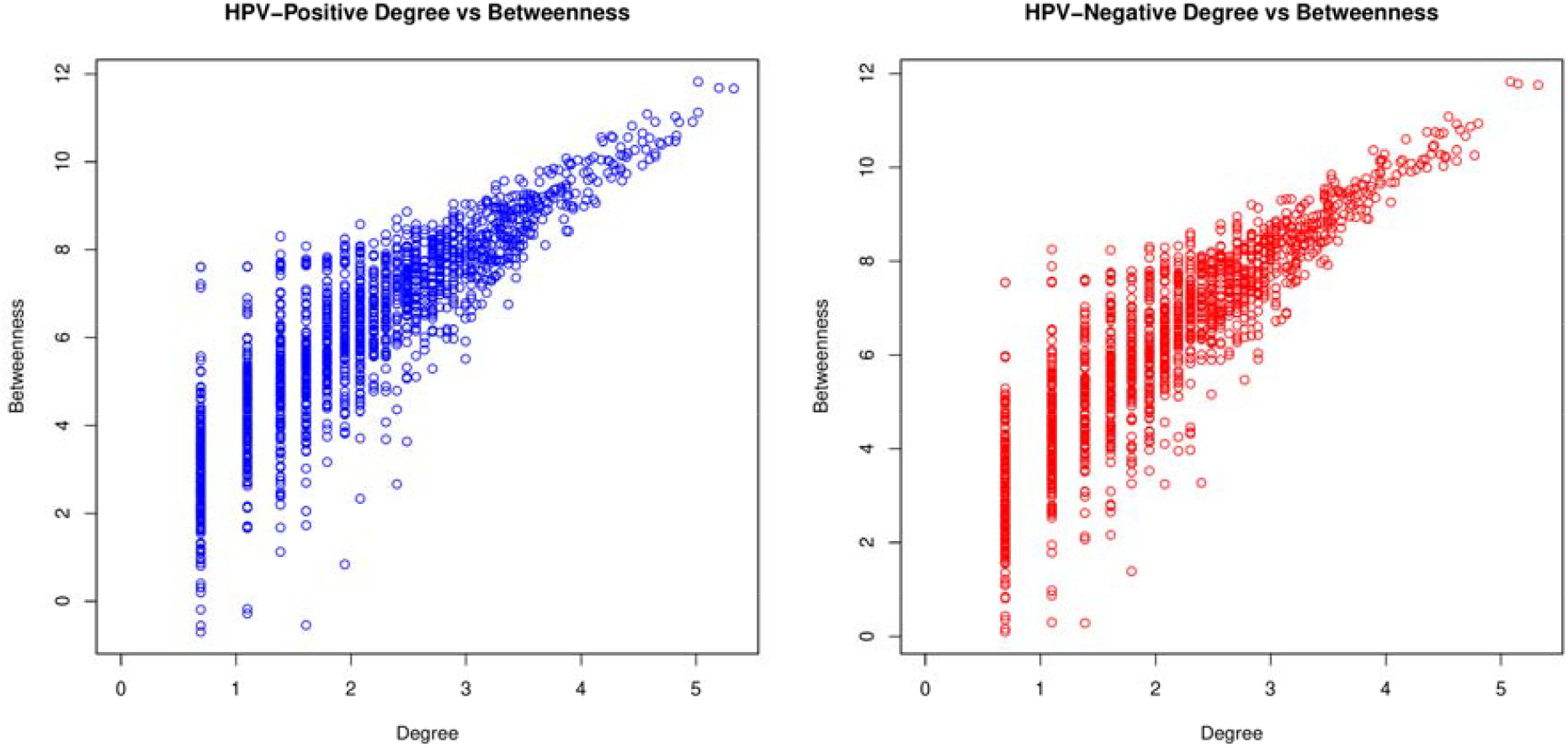
Degree-betweenness correlation plot for HPV-positive (left-side) and HPV-negative (right-side) HNSCC networks.

**Fig. 3:**
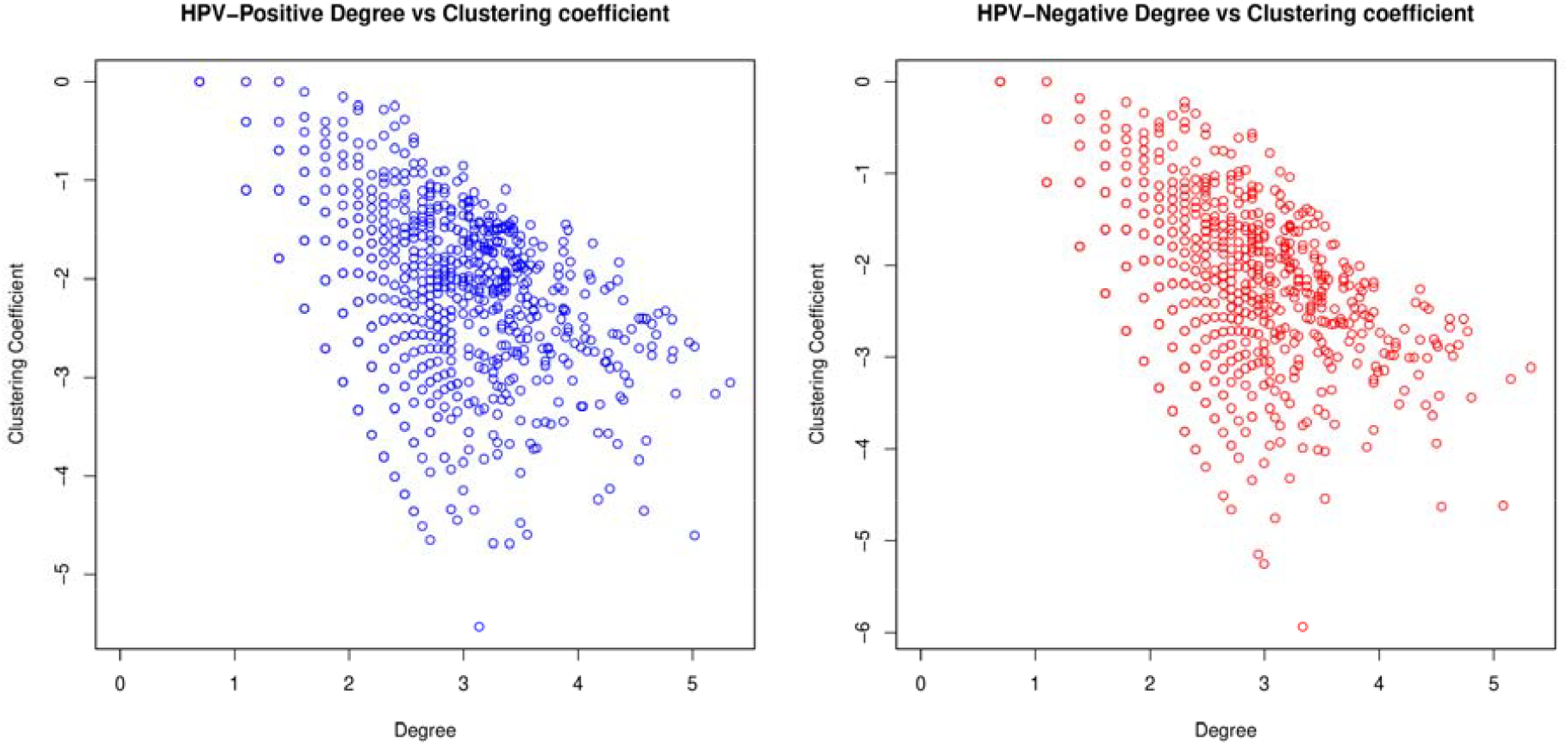
Degree-clustering coefficient correlation plot for HPV-positive (left-side) and HPV-negative (right-side) HNSCC networks.

### 3.2 Functional enrichment analysis of proteins present in HPV-positive and HPV-negative HNSCC

GO enrichment analysis disclosed that proteins present in the HPV-positive HNSCC network were involved in protein binding, cytoplasm, cell periphery, response to stimulus, anatomical structure development, multicellular organismal process, developmental process, positive regulation of biological process, positive regulation of cellular process, and cellular response to stimulus. Whereas, in HPV-negative HNSCC, proteins were mainly enriched in protein binding, cytoplasm, anatomical structure morphogenesis, cell periphery, response to stimulus, extracellular region, multicellular organismal process, tissue development, and developmental process (Fig. 4, Supplementary Table S1). The KEGG pathway analysis of HPV-positive HNSCC proteins were mainly involved in cell cycle, DNA replication, ECM-receptor interaction, protein digestion and absorption, small cell lung cancer, cell adhesion molecules, Human papilloma virus infection, cytokine-cytokine receptor interaction, AGE-RAGE signaling pathway in diabetic complications, and primary immunodeficiency; while, HPV-negative HNSCC proteins were mainly enriched in focal adhesion, ECM-receptor interaction, protein digestion and absorption, dilated cardiomyopathy, hypertrophic cardiomyopathy, pathways in cancer, arrhythmogenic right ventricular cardiomyopathy, small cell lung cancer, motor proteins, and Human papilloma virus infection (Fig. 5, Supplementary Table S2).

**Fig. 4:**
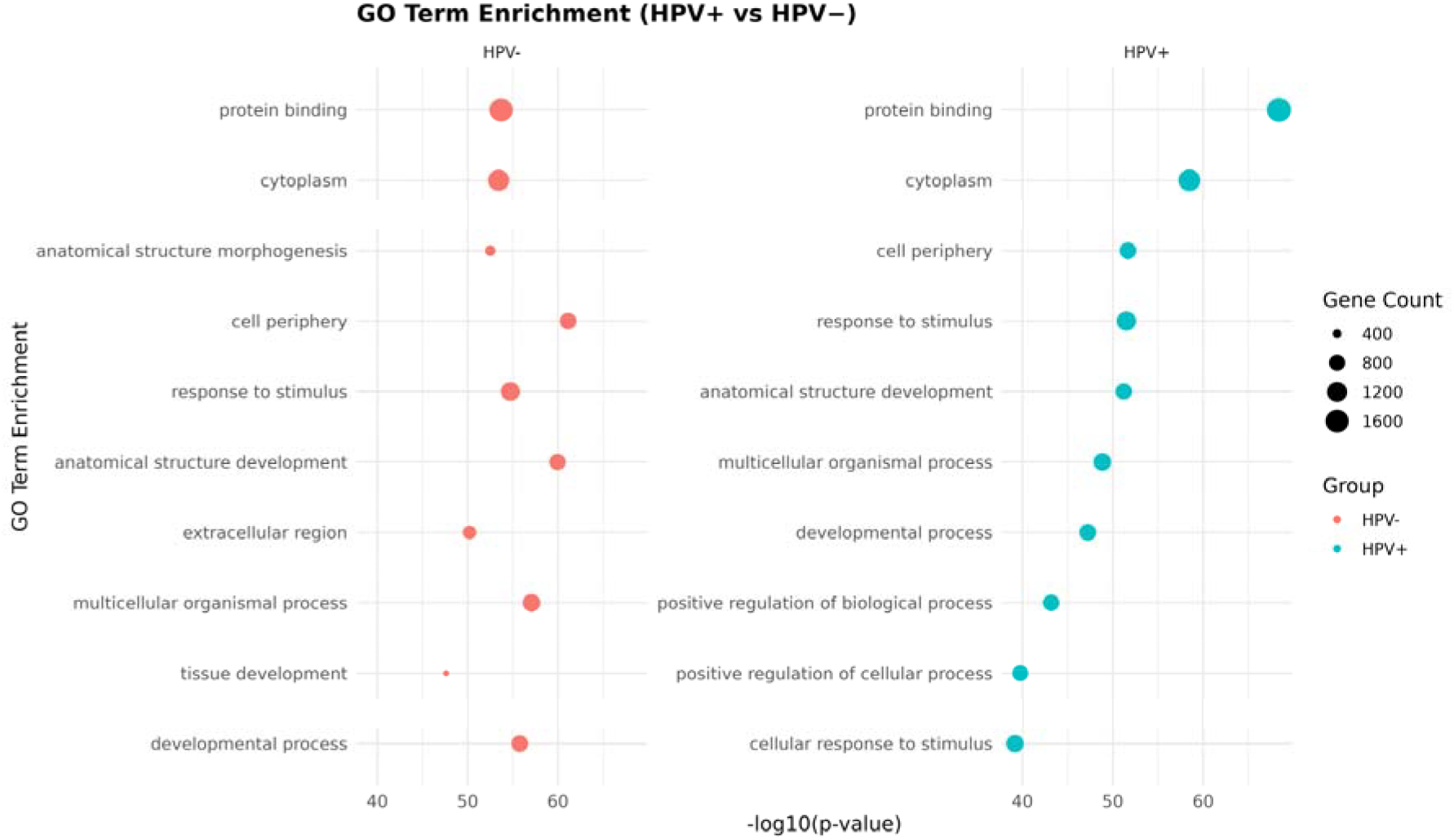
GO analysis of HPV-positive and HPV-negative proteins in HNSCC network.

**Fig. 5:**
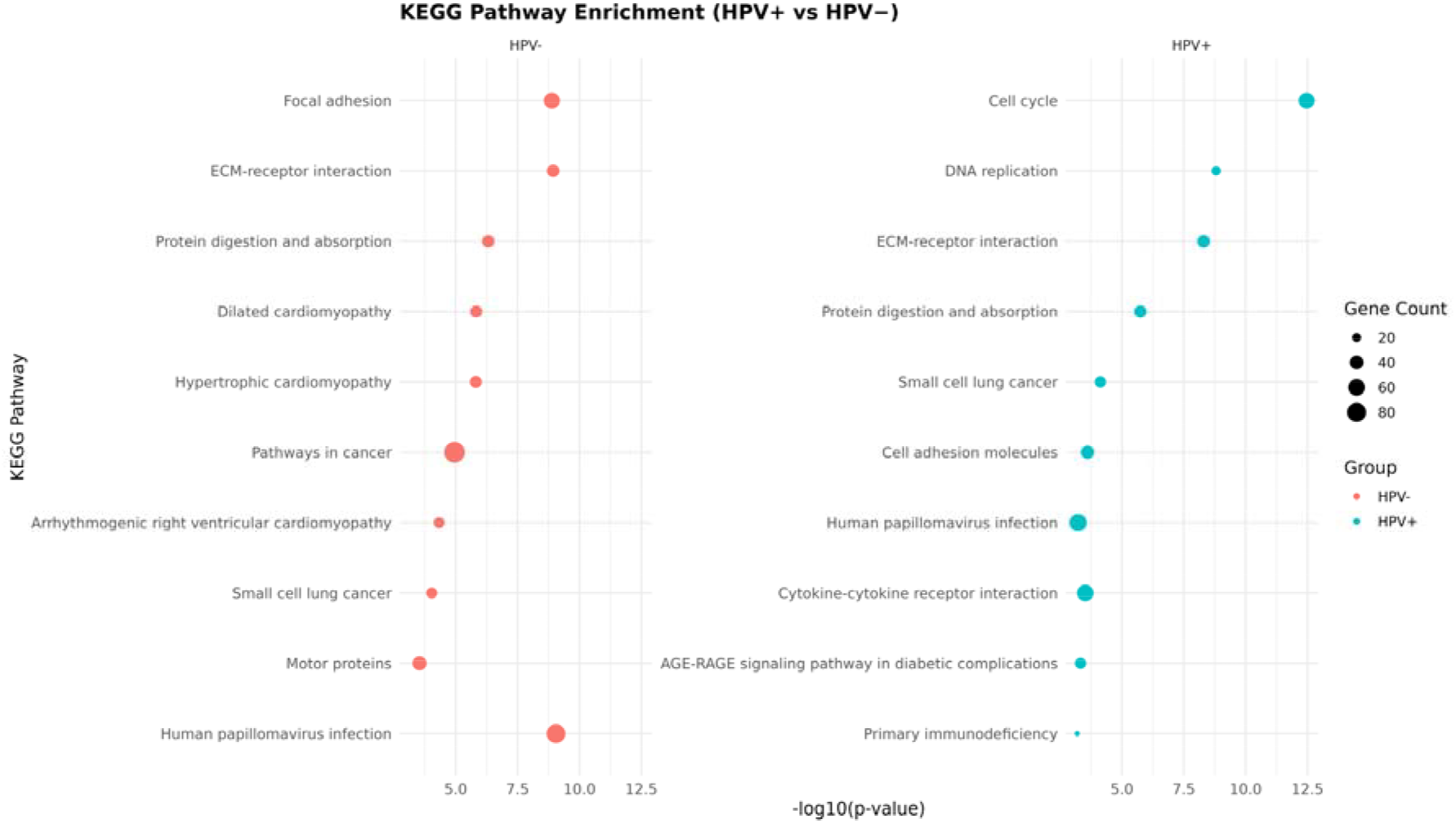
KEGG analysis of HPV-positive and HPV-negative proteins in HNSCC network.

### 3.3 Network vulnerability analysis

There were 286 vulnerable proteins in the HPV-positive HNSCC network, and 259 vulnerable proteins in the HPV-negative HNSCC network were identified from a one-node deletion approach. The two-node deletion approach revealed 1063 and 1081 vulnerable protein pairs in HPV-positive and HPV-negative HNSCC networks, respectively. Whereas, the three-node deletion approach revealed 38 and 302 vulnerable protein triplets in HPV-positive and HPV-negative HNSCC networks, respectively. Among them, 27 unique vulnerable proteins in HPV-positive (Supplementary Table S3) and 72 unique vulnerable proteins in HPV-negative (Supplementary Table S4) from one-node, two-node, and three-node deletion approaches.

### 3.4 Mutation analysis of vulnerable proteins

A total of 23 HPV-positive (Supplementary Table S5) and 70 HPV-negative (Supplementary Table S6) HNSCC vulnerable proteins having somatic mutation data were retrieved. After discarding the intron and silent mutation data, there were 20 and 62 HPV-positive and HPV-negative HNSCC somatic mutation data obtained, respectively. Similarly, 26 HPV-positive (Supplementary Table S7) and 67 HPV-negative (Supplementary Table S8) HNSCC vulnerable proteins with copy number data were filtered out. Among them, 01 and 17 vulnerable proteins were taken out in HPV-positive and HPV-negative HNSCC. To confirm the vulnerable proteins were functionally defective, proteins in combination with downregulated and copy-number variation or down-regulated and somatic mutation were taken out for further study.

### 3.5 Synthetic lethality and therapeutic target prediction

Using network-based and omics approaches, we identified three synthetic lethal partners for HPV-positive HNSCC vulnerable proteins and seven synthetic lethal partners for HPV-negative HNSCC vulnerable proteins to better understand their tumor-specific vulnerabilities. Three druggable targets were found for HPV-positive HNSCC vulnerable proteins (Table 1). Similarly, among the seven synthetic lethal partners for HPV-negative HNSCC vulnerable proteins, three proteins have known molecular inhibitors; three have not been reported yet, while one has an agonist (Table 1).

**Table 1:**
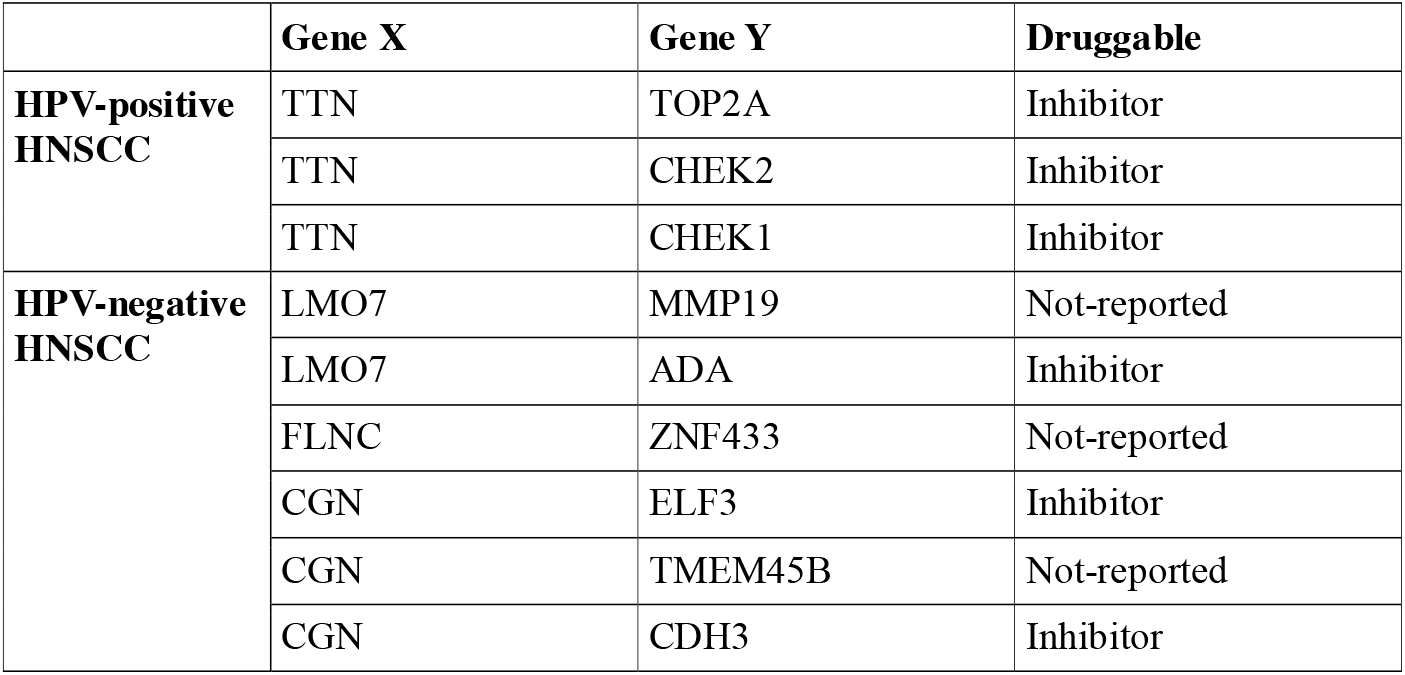
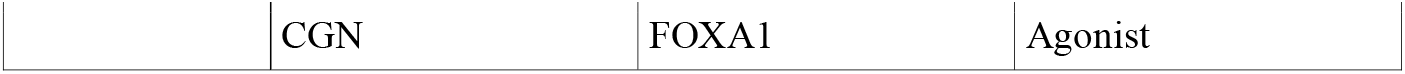
Synthetic lethal partners of vulnerable proteins with their druggable targets.

|  | Gene X | Gene Y | Druggable |
| --- | --- | --- | --- |
| **HPV-positive HNSCC** | TTN | TOP2A | Inhibitor |
|  | TTN | CHEK2 | Inhibitor |
|  | TTN | CHEK1 | Inhibitor |
| **HPV-negative HNSCC** | LMO7 | MMP19 | Not-reported |
|  | LMO7 | ADA | Inhibitor |
|  | FLNC | ZNF433 | Not-reported |
|  | CGN | ELF3 | Inhibitor |
|  | CGN | TMEM45B | Not-reported |
|  | CGN | CDH3 | Inhibitor |
|  | CGN | FOXA1 | Agonist |

## 4. Discussion

HNSCC is one of the leading cancers in the world and there is a requirement for a system-level approach to better understand the molecular mechanisms of HPV-positive and HPV-negative HNSCC. Analysis of PPI networks is useful to understand the underlying molecular mechanisms to improve the overall survival of patients. However, a combination of network-based and omics-based approaches is the most effective and well-suited for the identification of vulnerable proteins responsible for tumorigenesis in HPV-positive and HPV-negative HNSCC.

To explore the pathogenesis of HNSCC, we constructed the PPI networks for HPV-positive and HPV-negative HNSCC. The degree distribution, degree-betweenness correlation, and degree-clustering coefficient correlation of these networks were examined. Degree distribution plots of HPV-positive and HPV-negative HNSCC networks suggest that these networks follow a scale-free (power-law) topology, which is a key feature of a real biological system. Scale-free topology defines that most nodes have a minimum degree, and fewer nodes have a maximum degree. The degree-clustering coefficient correlation plot of HPV-positive and HPV-negative HNSCC networks indicated that there is a negative correlation between degree and clustering coefficient. We also analyzed the degree-betweenness correlation of the HPV-positive and HPV-negative HNSCC networks dataset, and the result showed that there is an overall positive correlation in the network.

To understand the role of proteins present in the HPV-positive and HPV-negative HNSCC networks, GO and KEGG pathway enrichment analyses were performed. The pathway analyses showed that the proteins of HPV-positive can adjust the cell cycle to influence HNSCC progression. While the proteins of HPV-negative HNSCC were mainly involved in pathways of cancer. Similarly, GO analyses showed that common enrichment terms such as protein binding, cytoplasm, and cell periphery were present in both HPV-positive and HPV-negative HNSCC. Moreover, HPV-positive HNSCC proteins also suggest active signaling and adaptive responses, likely reflective of the viral modulation of host cell machinery. In contrast, HPV-negative terms showed a stronger contribution of tissue remodelling and microenvironmental interactions.

Some proteins participating in the PPI network have more importance in terms of their position in the network, along with molecular functions, as compared to other participating proteins. Herein, 27 unique vulnerable proteins and 72 unique vulnerable proteins were identified in HPV-positive and HPV-negative HNSCC networks, respectively. Along with that, vulnerable proteins which has either downregulated as well as copy-number variation or downregulated as well as somatic variation were considered as functionally defective proteins. These proteins were defined as vulnerable as well as functionally defective proteins for HPV-positive and HPV-negative HNSCC networks because they represent loss-of-function or compromised-function events in cancer cells, making the cell dependent on backup pathways or compensatory genes, which can be targeted using synthetic lethality. Many targeted therapies aim to target nuclear receptor proteins or kinase domain-containing proteins by directly inhibiting the activated protein product (Benstead-Hume *et al*., 2019). However, such approaches are ineffective to repair tumor suppressor genes or their protein products, especially when these are inactivated by a truncating mutations (Khoo *et al*., 2014). To overcome this limitation, the concept of synthetic lethality is gaining considerable interest. Herein, a total of 07 HPV-positive and 16 HPV-negative vulnerable proteins were found, which were also functionally defective. Among them, TOP2A, CHEK2, and CHEK1 were identified as SL partners of TTN in the HPV-positive HNSCC. In the HPV-negative HNSCC, ADA and MMP19 were identified as SL partners of LMO7; TMEM45B, FOXA1, CDH3, and ELF3 genes were identified as SL partners of CGN; and ZNF433 was identified as an SL partner of FLNC. TTN is one of the most commonly mutated genes in various cancers, such as colon cancer (Baumert *et al*., 2022) and HNSCC (Shaikh *et al*., 2019). It has been well reported that a mutation in the TTN gene may lead to abnormal myosin function, which results in abnormal muscle fibre growth (Savarese *et al*., 2018). Whereas reports suggest that high expression of the LMO7 gene was associated with better overall survival in oropharyngeal squamous cell carcinoma (Israelsson *et al*., 2024). The LMO7 gene is involved in various functions such as cell migration, stabilizing role in adherens junctions, and acts as a transcription factor of muscle-related genes (Miyoshi *et al*., 2008; Hu *et al*., 2011; Holaska *et al*., 2006). Paschoud et al. suggested that CGN is upregulated in lung adenocarcinoma, but its expression is reduced in squamous cell carcinoma (Paschoud *et al*., 2007). Moreover, Ai et al. suggested that FLNC facilitates the migratory capacity of tumor cells in prostate cancer (Ai *et al*., 2017). Thus, from the above finding, we can say that vulnerable proteins are mainly involved in cytoskeleton organization, muscle contraction, and cellular architecture.

To explore the therapeutic relevance of the identified vulnerable proteins in HPV-positive and HPV-negative HNSCC, TOP2A, CHEK2, and CHEK1 in HPV-positive HNSCC have known pharmacological inhibitors. These synthetic lethal partners are associated with known inhibitors, which are clinically approved. Similarly, ADA, CDH3, and ELF3 in HPV-negative HNSCC also associated with pharmacological inhibitors, but ADA has clinically approved inhibitors, while CDH3 and ELF3 do not have clinically approved inhibitors. However, inhibitors targeting TMEM45B, MMP19, and ZNF433 have not yet been reported. Therefore, these genes and their associated synthetic lethal interactions provide promising avenues for future therapeutic exploration in HNSCC.

## 5. Conclusion

Our findings revealed the most vulnerable proteins, viz., TTN, CGN, LMO7, and FLNC in HPV-positive and HPV-negative HNSCC. These proteins play a critical role in maintaining network integrity, as their removal leads to the network collapse. Moreover, they were found functionally defective, harboring both CNV and SNV. The functional enrichment analysis suggested that the cytoskeleton organization and muscle contraction pathways may have contribution in the development of HPV-positive and HPV-negative HNSCC. Collectively, these findings highlight the potential of vulnerable proteins as novel therapeutic targets for future drug development.

## Data availability

The supplementary materials can be found online with this article.

## Acknowledgments

Vaibhav Vindal would like to thank the Anusandhan National Research Foundation (ANRF/PAIR/2025/000012/PAIR-A), Indian Council of Medical Research (ICMR), New Delhi (ISRM/12(72)/2020, ID: 2020-2951), and Department of Biotechnology (DBT), Government of India (No. BUILDER-DBT-BT/INF/22/SP41176/2020), for their financial support to Lab. Further, Swapnil Kumar would like to acknowledge the ICMR for providing financial assistance as the Senior Research Fellowship (Grant No. 3/2/2/113/2019/NCD-III, ID: 2019-6723) and the IoE-UoH for the Publication based incentives. Avantika Agrawal would also like to acknowledge the IoE-UoH for the Publication-based incentives.

## CRediT authorship contribution statement

Conceptualization: Avantika Agrawal, Swapnil Kumar, and Vaibhav Vindal; Methodology: Avantika Agrawal and Swapnil Kumar; Formal analysis and investigation: Avantika Agrawal; Writing - original draft preparation: Avantika Agrawal, Writing - review and editing: Avantika Agrawal, Swapnil Kumar, and Vaibhav Vindal; Supervision: Vaibhav Vindal.

## Supplementary Figure Legends

**Supplementary Figure F1: PPI network of HPV-positive HNSCC**.

**Supplementary Figure F2: PPI network of HPV-negative HNSCC**.

## Supplementary Table Legends

**Supplementary Table S1: Significantly enriched MF, BP, CC terms associated with vulnerable proteins in HPV-positive and HPV-negative HNSCC networks**.

**Supplementary Table S2: Significantly enriched KEGG terms associated with vulnerable proteins in HPV-positive and HPV-negative HNSCC networks**.

**Supplementary Table S3: Common proteins in one-node, two-node pairs, and three-node triplets in the HPV-positive HNSCC network**.

**Supplementary Table S4: Common proteins in one-node, two-node pairs, and three-node triplets in the HPV-negative HNSCC network**.

**Supplementary Table S5: Somatic mutation in the HPV-positive vulnerable proteins HNSCC**.

**Supplementary Table S6: Somatic mutation in the HPV-negative vulnerable proteins HNSCC**.

**Supplementary Table S7: Copy number variation in the HPV-positive vulnerable proteins HNSCC**.

**Supplementary Table S8: Copy number variation in the HPV-negative vulnerable proteins HNSCC**.

